# The cellular states and fates of shed intestinal cells

**DOI:** 10.1101/2022.10.03.510592

**Authors:** Keren Bahar Halpern, Yael Korem Kohanim, Adi Biram, Adi Egozi, Ziv Shulman, Shalev Itzkovitz

## Abstract

The intestinal epithelium is replaced every few days^1^. Enterocytes are shed into the gut lumen predominantly from the tips of villi^3,4^, and are believed to rapidly die upon their dissociation from the tissue. However, technical limitations prohibited studying the cellular states and fates of shed intestinal cells. Here, we used bulk and single cell RNA sequencing of mouse intestinal fecal washes to demonstrate that shed epithelial cells remain viable and up-regulate distinct anti-microbial programs upon shedding. We further identify abundant shedding of immune cells, a process that is elevated in DSS-induced colitis. We find that fecal host transcriptomics mirrors changes in the intestinal tissue following perturbations. Our study suggests potential functions of shed cells in the intestinal lumen and demonstrates that host cell transcriptomes in intestinal washes can be used to probe tissue states.

## Introduction

The gut epithelium is one of the cell populations with the highest turnover rates in mammals. The mouse epithelium is replaced every 3-5 days^1^ and the human gut sheds more than 30 grams of cells every day^2^. Gut cells are shed into the lumen predominantly from the tips of villi in the small intestine and from the inter-crypt epithelium in the large intestine^3,4^. This shedding is a result of replacement by cells that are continuously produced at the crypts and driven by mitotic pressure^5-7^. It is believed that shed cells enter anoikis, a process in which the cell loses contact with adjacent cells as well as with the extracellular matrix^8–12^. Notably, some studies showed shedding of cells without apoptotic markers in the human colon^13^, suggesting that shed cells may continue to be viable and perhaps even functional. This hypothesis has thus far not been systematically examined. Understanding the biology of intestinal shed cells has important clinical implications. Several studies analyzed stool host transcriptomes to investigate gut development^14^, disease progression^15^ or impacts of therapeutic regimens^16^. Fecal wash host transcriptomes collected during colonoscopies have been shown to be statistically powerful classifiers of histological remission in patients with Inflammatory Bowel Disease (IBD)^17^. These findings highlighted the potential prognostic value of shed cell transcriptomics in the gut. However, it remains unclear whether these informative fecal transcriptomes are remnants of cells that have died immediately after shedding, or whether shed cells may still be viable and even functionally distinct upon their dissociation from the tissue.

Here, we present a comprehensive analysis of mouse fecal-wash host-transcriptomes throughout the mouse intestinal tract. We performed bulk and single cell RNA sequencing (scRNAseq) as well as single molecule FISH (smFISH) and intra-vital imaging to study the identities and cellular states of shed cells. We show that substantial amounts of immune cells are regularly shed during homeostasis, and that immune cell shedding increases in DSS-induced colitis. Our work suggests that shed cells may continue to carry functions in the lumen, and establishes the power of fecal wash host transcriptomics in mirroring mucosal physiology and pathology.

## Results

### Fecal wash host transcriptomes have distinct segment-dependent profiles

To study the molecular identities of shed intestinal cells in mice, we divided the gut into 5 segments, 3 equal-length segments spanning the small intestine (duodenum, jejunum and ileum) and 2 equal-length segments spanning the large intestine (proximal and distal colon, Fig. 1a). We flushed each segment with PBS into a tube (i.e. Fecal wash). In addition, we collected a small piece of tissue into a separate tube. We performed bulk RNA sequencing using the mcSCRBseq protocol^18^ (Supplementary Table 1). Principal Component Analysis (PCA, Fig. 1b) of the transcriptomic data revealed two axes of variations. The first PC correlated with the intestinal segments, highlighting large variation between the small and large intestine and smaller variations among sub-segments (Fig. 1b). The second PC separated the tissue and fecal wash samples.

**Fig. 1.**
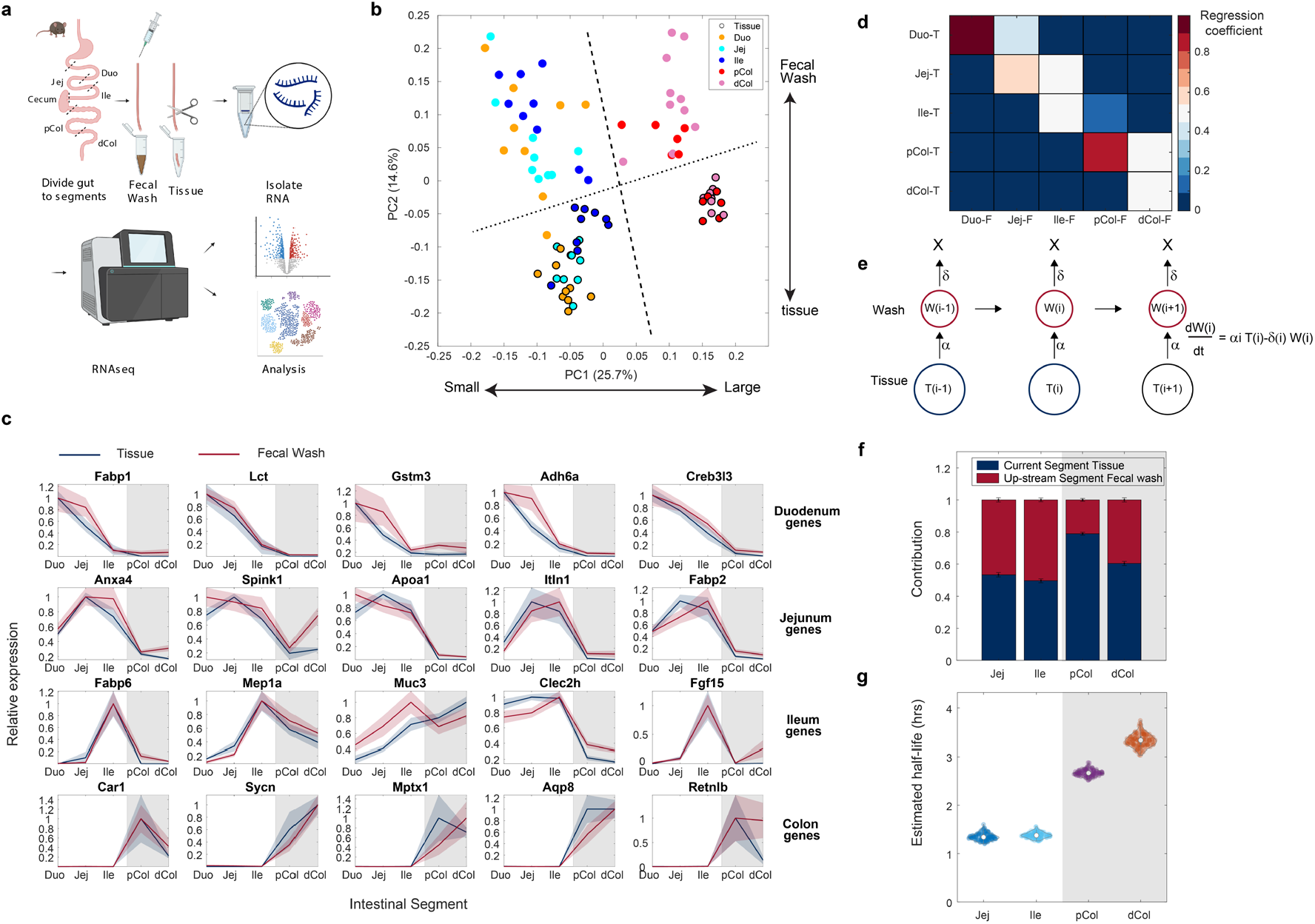
Fecal wash host transcriptomes have distinct segment-dependent profiles. **a**. Illustration of fecal wash and tissue sample collection and experimental layout. Duo - Duodenum, Jej – Jejunum, Ile – Ileum, pCol-Proximal colon, dCol - Distal colon. Created with BioRender.com. **b**. Principal component analysis (PCA) of all control samples. Sample are color coded by intestinal segment source. Black circles denote tissue samples. Numbers in parentheses show percent of explained variance by each PC. **c**. Gene expression segmental profiles of tissue samples (blue) or fecal wash samples (red) along the gastrointestinal tract. Gray shades denote the colon. **d**. Multi-linear regression of the fecal wash transcriptomes based on the segmental tissue transcriptomes. Shown are the regression coefficients. **e**. Mathematical model of shed cell turnover. Intestinal segment correlates with time, shed cells are produced at rate α and removed at rate δ. **f**. Fecal wash transcriptome predominantly constituted a weighted average of the matching tissue contribution (blue) and previous segment fecal wash (red, Methods). **g**. Estimated half-life of shed cells throughout the gastrointestinal tract (Methods). Data was bootstrapped to obtain statistics, circles denote the medians, vertical bars denote the Inter-Quartile Range (IQR).

Variations among segments in fecal wash samples from the small intestine were less pronounced compared to the tissue samples (between segment /within segment ratio of 1.18 for tissues vs. 1.13 for fecal washes, p< 3e-196). We reasoned that fecal washes at each segment might still contain transcripts from cells that were shed in up-stream segments. Genes with clear segmentally varying profiles, such as the proximal Fabp1 and the distal Fabp6^19^ showed broadly similar segmental profiles in the fecal washes (Fig. 1c). Notably, fecal wash segmental profiles often declined less abruptly compared to the tissues (e.g. Gstm3 and Clec2h, Fig. 1c), or even exhibited a shift in the segmental peak (e.g. Itln1 and Fabp2, Fig. 1c), consistent with a picture of elongated lifetime of the shed cells or their mRNA content.

Using multi-linear regression (Fig. 1d), we found that fecal wash transcriptomes predominantly constituted weighted averages of the transcriptomes of the matching and immediate up-stream tissue segments. The proximal colon fecal wash (pCol-F) had lower representation of the upstream ileum samples, potentially indicating increased degradation of mRNA at the cecum. Since the intestinal segment correlates with transit time, we formulated a mathematical model of cell shedding (Methods, Fig. 1e-f) that facilitates inferences of shed cell transcriptome lifetimes. The model incorporated the contribution of shed cell transcriptomes at each segment (shedding rate α) and the luminal transcript degradation rate (δ). Estimated mRNA lifetimes were on the order of 1.2-1.6 hours in the small intestine and 2.5-3.7 hours in the colon (Fig. 1g), assuming a total transit time of 8 hrs throughout the mouse gastrointestinal tract with faster transit in the small intestine compared to the large intestine^20–22^ (Methods). Shed cells, or their mRNA content therefore seem to survive for several hours in the gut lumen.

### Differences in gene expression between shed and tissue cells

To explore the differences in expression between the tissue and the shed cells, we performed differential gene expression analysis (Fig. 2, Extended Data Fig. 1, and Supplementary Table 2). Genes that were upregulated in fecal washes included immune-related genes such as Adrgre1 encoding the macrophage marker F4/80, the T-cell marker gene Ccl5, as well as bacterial defense genes such as Nfkbia and Cdkn1a (Fig. 2a). Genes enriched in the tissue included the proximal and distal small intestinal genes Fabp1 and Fabp6^19^ respectively, as well as mesenchymal genes such as Col1a1 (Fig. 2a). Gene Set Enrichment Analysis (GSEA)^23^ analysis indicated that fecal washes from both small and large intestine were further enriched in genes associated with TNFα signaling, IFNγ/α signaling, and hypoxia pathways, and depleted in pathways such as oxidative phosphorylation and cell cycle pathways G2M checkpoint and E2F targets (Fig. 2b). In addition, many villus tip markers were elevated in fecal washes, including Fos, Ada and Nt5e (Fig. 2a), indicating an increase cell shedding from the villus tip^24^. To infer cell type composition of both fecal washes and tissues we used the cellanneal computational deconvolution software^25^. We found that fecal washes were enriched in villus tip cells. Notably, fecal washes were also enriched in neutrophils, monocytes and macrophages (Fig. 2c,d).

**Fig. 2.**
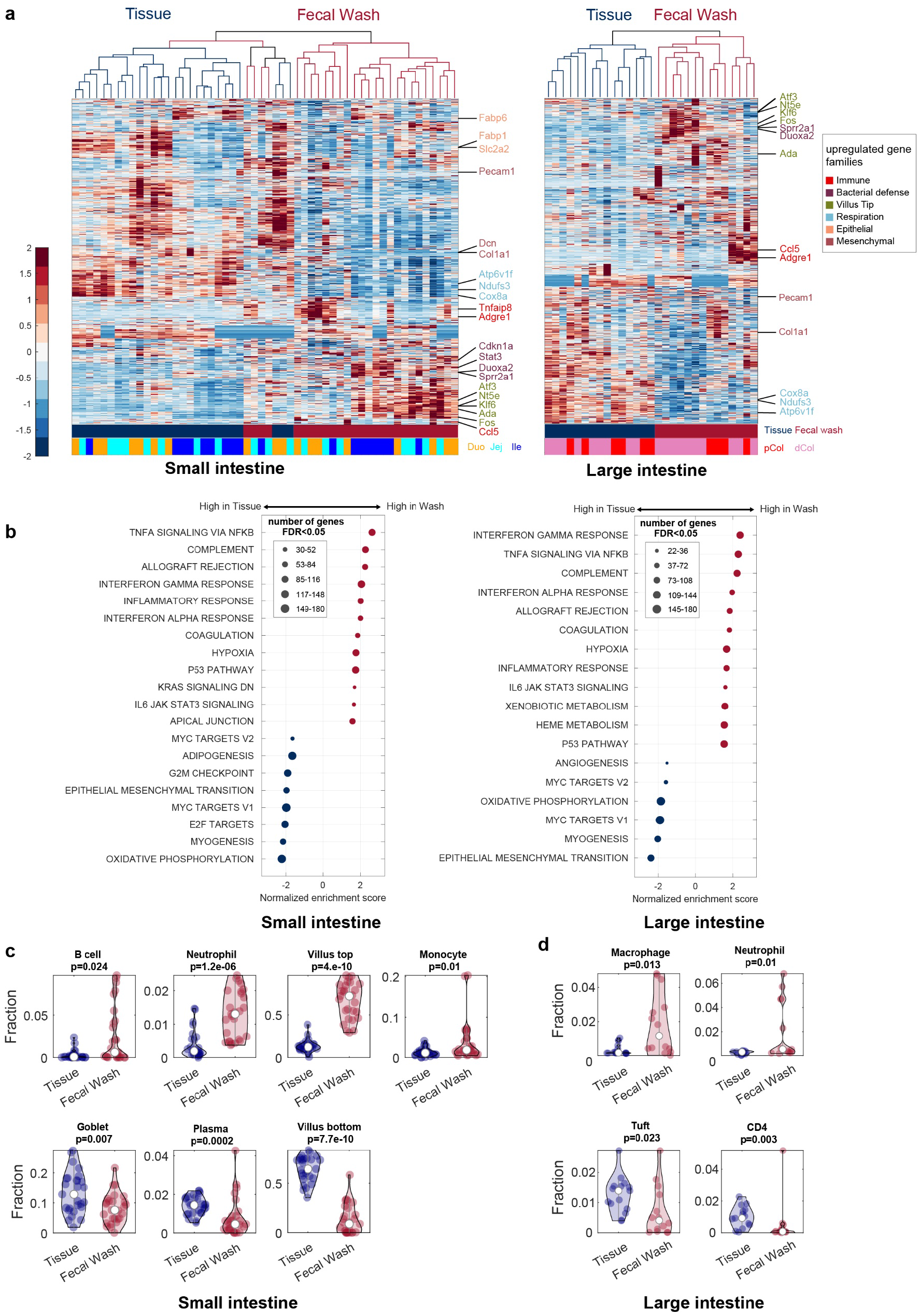
Differences in gene expression between the tissue and the shed cells. **a**. Hierarchical clustering of fecal wash samples (red branches) or Tissues (blue branches). Left, small intestine samples, right, large intestine samples. Samples naming nomenclature and color coding; Duo – Duodenum - Orange, Jej – Jejunum - Cyan, Ile – Ileum - Blue, pCol-Proximal colon - Red, dCol - Distal colon - Magenta. Representative genes are shown on the right. Expression values are Z scores (methods). **b**. GSEA over the Hallmark gene sets. Left - small intestine, right - large intestine, included are gene sets with q-value <0.05. Gene sets elevated in fecal washes (red circles) were associated with immune cell pathways, while gene sets elevated in the tissue samples (blue circles) included cell proliferation and respiratory pathways. FDR, false discovery rate; **c-d**. Estimated fractions of different cell types in fecal washes and tissue samples inferred by computational deconvolution^25^. **c** – small intestine, **d** – large intestine. White circles are medians, vertical bars denote the Inter-Quartile Range (IQR). P-values computed using Kruskal-Wallis tests.

### Fecal washes contain diverse intact viable cells

The inferred abundance of immune cells in fecal washes was unexpected, as cell shedding is thought to be dominated by epithelial cells in homeostasis^7^. To further establish this finding, we sorted fecal wash cells into 384 well capture plates and sequenced them using the MARSseq protocol (Fig. 3a, Extended Data Fig. 2)^26^. Consistent with the computational deconvolution results, our scRNAseq atlas included abundant monocytes, B-cells, Cycling monocytes, Mast cells and T-cells, in addition to epithelial Enterocytes, Goblet cells and Tuft cells (Fig. 3a,b). Using smFISH of fecal wash cells, we validated the existence of intact enterocytes (marked by Ada), B-cells (marked by Cd19) and monocytes (marked by Lyz1, Fig. 3c-e). Notably, shed cells exhibited bright transcription sites, indicating active transcription (Fig. 3c, white arrow). Using intravital imaging (Methods), we identified monocyte shedding in a CX3CR1^GFP^ mouse model (Fig. 3f, supplementary movie 1, Methods). Shed cells therefore seem to be diverse, viable and transcriptionally active.

**Fig. 3.**
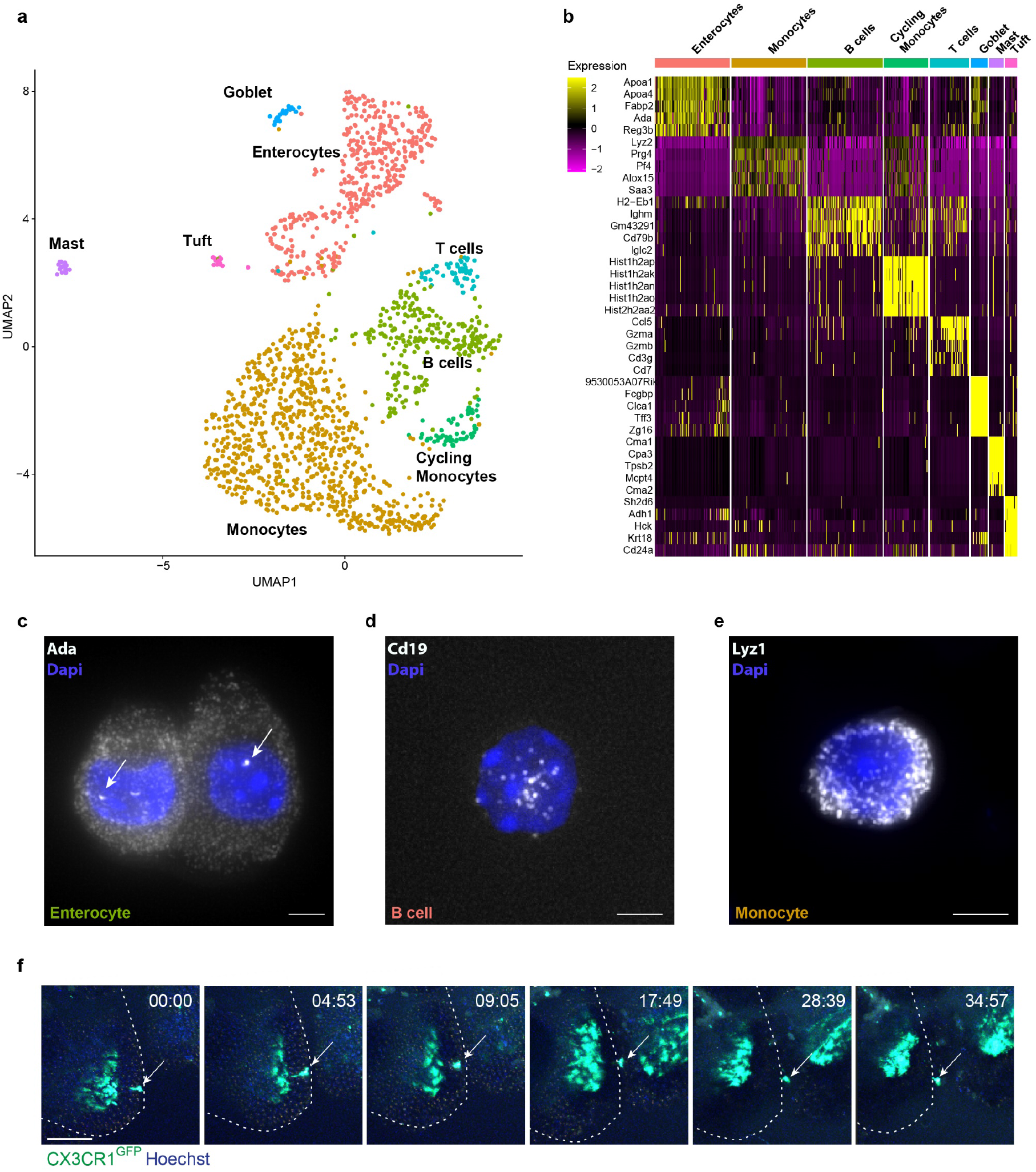
single cell analysis of fecal washes reveals diverse cell type presence. **a**. UMAP (Uniform Manifold Approximation and Projection) visualization of single cells obtained from small intestine fecal washes analyzed by single cell MARSseq. Cells clustered into 8 groups colored by cell type annotation. n=1,717 cells from 4 mice. **b**. Heatmap showing the top 5 differentially expressed genes in each cell type. Expression is log-normalized by Seurat^27^, cells randomly down-sampled to a maximum of 100 per cell type. **c-e**. smFISH validation of selected cell type markers found in **b**, Ada mark enterocytes, white arrow marks an active transcription site, Cd19 marks B-cell and Lyz1 marks monocytes. Scale bar is 5μm. **f**. Intra-vital imaging of CX3CR1^GFP^ mice injected with 10μg TNF-α intraperitoneally in PBS 20 min prior to the surgery (Methods). Capture movie was 35 minutes. Scale bar is 70μm. Dashed lines mark villus border. Arrow highlights a shedding monocyte (supplementary movie 1).

### Shed enterocytes upregulate anti-inflammatory modules

Our findings that shed cells continue to be viable and transcriptionally active, suggested that they might carry luminal functions that are distinct from their tissue roles. In order to understand the changes enterocytes undergo after shedding, we next combined our fecal wash scRNAseq measurements with scRNAseq of tissue enterocytes^28^ (Fig. 4a). We used landmark genes^24^, to infer the location of each cell along the crypt-villus axis, denoted as ‘zonation score’, ranging from 0 at the crypt to 1 at the villus tip (Fig. 4a). We found that the fecal wash cells had a clear villus tip signature (median zonation score 0.73 for shed enterocytes and 0.16 for tissue enterocytes, p=8e-102, Methods, Fig. 4b). Notably, differential gene expression between the shed enterocytes and tissue enterocytes with matched zonation distributions identified significant expression changes (1,589 genes out of 7,021 genes with expression above 10^−5^ were significantly different between the groups at false discovery rate (FDR) of 0.1). Genes that were upregulated in fecal wash enterocytes compared to tissue enterocytes included the anti-microbial genes Reg3g and Reg3b, the redox-signaling genes Duoxa2 and Duox2 and the pro-inflammatory response genes Saa1 and Cd74 genes (Fig. 4c). Shed enterocytes were enriched in genes associated with TNFα signaling, IFNγ/α signaling, and hypoxia pathways in addition to IL6 signaling, and the JAK STAT pathway, and depleted in oxidative phosphorylation related genes (Fig. 4d). Our data indicates that shed enterocytes up-regulate anti-microbial and immune-related processes.

**Fig. 4.**
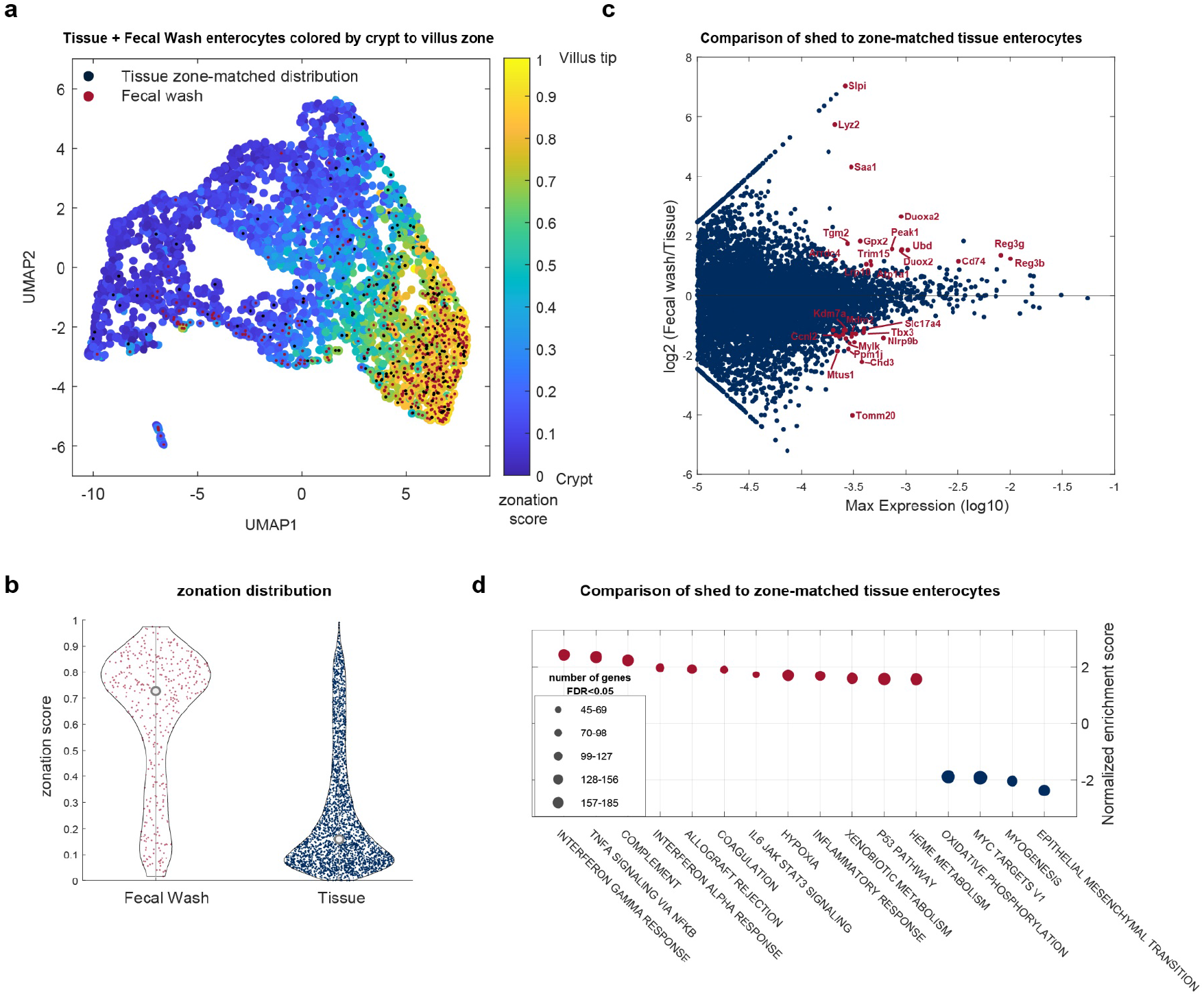
Enterocytes are predominantly shed from villi tips and elevate anti-microbial pathways. **a**. UMAP visualization of single enterocytes obtained from both fecal washes and jejunum tissues. Cells are colored by zonation score (Methods), which correlates with crypt-villus-tip axis location. Red dots mark fecal wash cells; Black dots mark tissue cells with distribution of zonal coordinates that matches that of shed cells zonal coordinates (Methods). **b**. zonation distribution of all fecal wash enterocytes (red dots) and tissue enterocytes (blue dots) present in **a**. Medians are marked with grey circles. **c**. MA plot of differential gene expression between fecal wash and matched distribution tissue cells. Each dot is a gene, x-axis is the log10 of max expression, y-axis is log2 ratio between mean expression in fecal wash enterocytes and tissue enterocytes, p-value is computed using Wilcoxon rank-sum tests. Names of representative up-regulated genes (q-values below 0.1) are marked in red (q-values computed using Benjamini-Hochberg FDR correction). **d**. GSEA over the Hallmark sets. Shown are all gene sets with q-value <0.05. Gene sets up regulated in shed enterocytes (red circles) included immune cell pathways, gene sets up regulated in tissue enterocytes (blue circles) included cell proliferation and respiratory pathways.

### Gut microbial alteration changes the shed cell composition

We next asked if fecal wash host mRNA composition changes under perturbations that modulate microbial loads and challenge barrier function. To this end, we applied two perturbations: long term broad-spectrum antibiotics (ABX) exposure and DSS (dextran sulfate sodium) induced colitis (Fig. 5a). We compared the induction of gene expression in colon post DSS treatment (Fig. 5b) in both tissue and fecal washes. We found that the changes in the fecal wash host transcriptomics mirrored the changes measured in the tissue (R spearman 0.29, p=3.1e-308). Elevated genes were consistent with ones previously reported to be induced in both mice and human colitis^29^ (median fold change 7.9 in tissue, 4.9 in fecal washes). These genes included S100a8 and S100a9, encoding the subunits of the calprotectin protein, a known marker of gut inflammation^30^. As well as the anti-microbial genes, Reg3g and Reg3b and immune cell markers Il1b and Cxcl2 (Extended Data Fig. 3a). ABX yielded a slightly lower, yet statistically significant correlation between the tissue and fecal wash transcriptomics (R spearman 0.11, p=1.7e-39, Fig. 5C). Changes including a strong decrease in the antimicrobial genes Reg3g and Reg3b in both tissue and fecal washes. Other downregulated genes were consistent with ones previously reported to be repressed in ABX treated mice^31^ (Extended Data Fig. 3b). These genes included Ang4, an antimicrobial factor involved in innate immunity^32^ as well as Retnlb encoding Relmβ a goblet cell secreted factor that selectively target gram negative bacteria in the gut^33^. Computational deconvolution indicated a significant increase in fecal wash immune cell fraction in DSS colitis and a decrease in ABX, both only seen in the colon (Fig. 5d,e, Supplementary Table 3). GSEA indicated that fecal washes from DSS mice compared to control mice were enriched in genes associated with Jak/Stat3 and Il2/Stat5 signaling, known pathways upregulated in colitis^34,35^. These pathways were not elevated in the ABX mice fecal washes (Supplementary Table 4). Thus, fecal wash host transcriptomics mirrors pathological changes in the intestinal tissue.

**Fig. 5.**
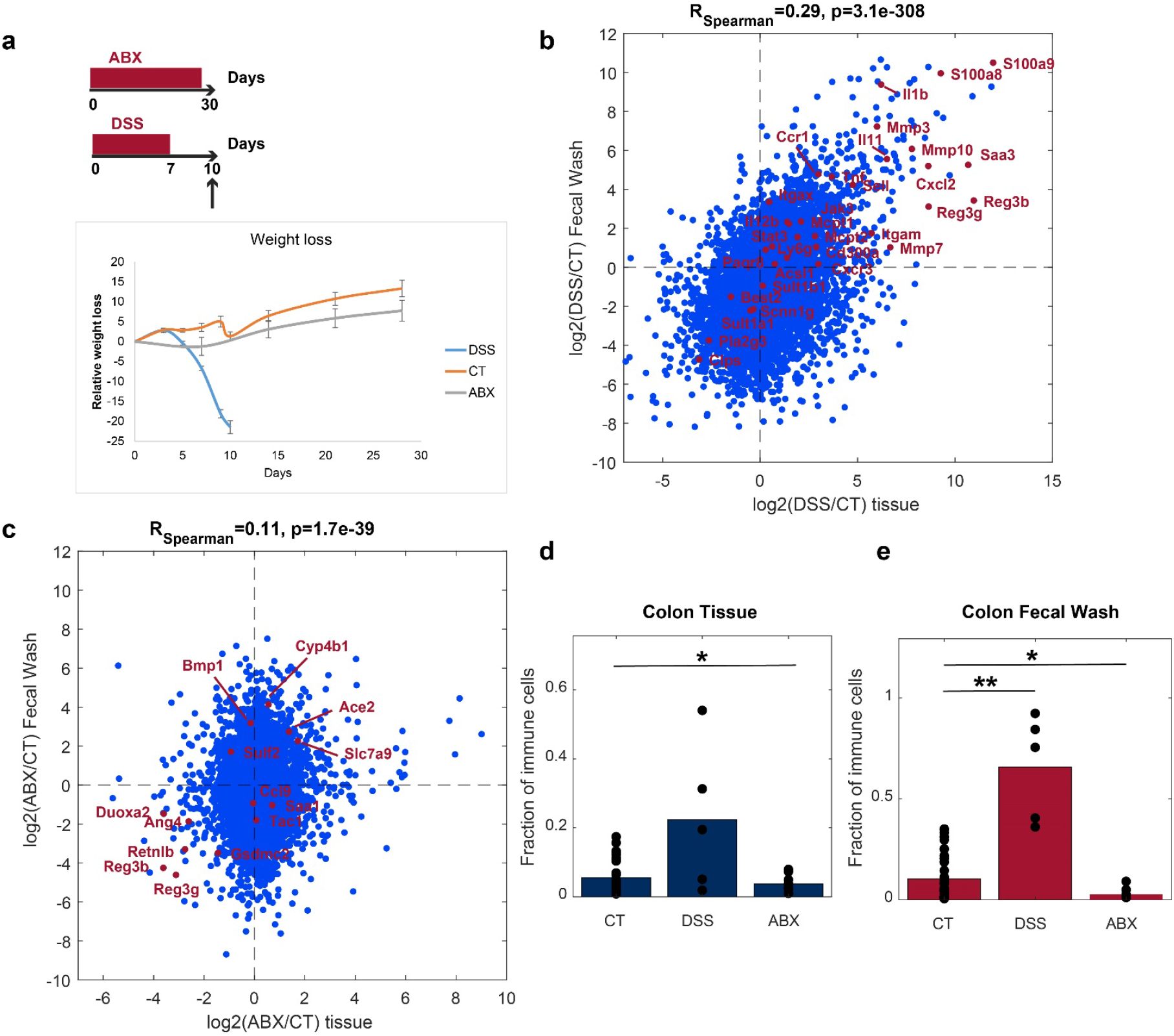
Fecal washes mirror perturbed tissue states. **a**. Time scale scheme of mice perturbations. ABX-antibiotics, DSS – dextran sulfate sodium induced colitis. Weight loss graph show changes in body weight throughout the drag administration. **b**. Scatter plot of log2 gene expression ratio between DSS and control mice in colon tissue (x-axis) or colon fecal washes (y-axis). Each dot is a gene. Red dots are inflammatory-related genes previously shown to be upregulated in colon tissues from DSS-colitis^29^. Spearman correlation 0.29, p-value = 3.1e^-308^. **c**. Scatter plot of log2 gene expression ratio between ABX and control mice in colon tissue (x-axis) or colon fecal washes (y-axis). Red dots are genes previously shown to change in colon tissues from after ABX gavage treatment^30^ Spearman correlation 0.11, p-value = 1.7e^-39^. **d-e**. Changes in fractions of immune cells between treatments calculated from computational-deconvolution cell compositions of colon tissue (**d**) or colon fecal wash (**e**). CT – control. *p-value<0.05 **p-value<0.01.

## Discussion

The traditional view of intestinal cell turnover posits that enterocytes die immediately upon their dissociation from the tissue. Contrary to this view, our study demonstrated that shed epithelial cells remain viable for a few hours in mice. Using single cell RNA sequencing and comparison to the zonated programs of tissue enterocytes (Fig. 4), we found that enterocytes are shed predominantly from the villus tip, but are transcriptionally distinct from their tissue counterparts. Enterocytes have been shown to elevate the expression of dozens of genes at the very tip of the villi, shortly before their shedding^24^. Our study demonstrates that they further up-regulate genes, including anti-microbial programs, after their shedding. Release of viable shed cells might serve to modulate the luminal microbial composition.

Our study demonstrated continuous shedding not only of epithelial cells, but also of diverse populations of immune cells. Trans-epithelial migration of neutrophils is a hallmark of gut inflammation^33^. Using scRNAseq, smFISH and intra-vital imaging, we showed that immune cells that include both myeloid and lymphoid cells are normally shed in un-perturbed mice as well. It will be important to explore the potential functions of these cells within the lumen. We also found that immune cell shedding was modulated by the microbiome, increasing in DSS-colitis and decreasing upon broad-spectrum antibiotic treatment. The elevated immune-cell shedding in DSS-colitis, a mouse model for IBD, are in line with recent studies in human, that demonstrated elevated shedding of immune cells during IBD flares^17^.

Our controlled measurements of transcriptomes of tissue and shed cells along the gastrointestinal tract enabled estimates of their turnover dynamics (Fig. 1d-g). We found that segmental fecal washes represent a mixture of cells shed from the corresponding intestinal segments, as well as from up-stream segments. The cecum seems to form a barrier that potentially stalls the cells, increases their rate of death or the degradation rate of their transcriptomes, as representation of ileal segments in washes of the proximal colon was lower (Fig. 1d,f). Notably, we estimate that shed cells survive longer in the colon compared to the small intestine (Fig. 1g). Although the colon is substantially shorter than the small intestine, transit time along this segment is significantly longer^20–22^, yet the distal colon had a substantial contribution from the proximal colon shed cells (Fig. 1f). The shorter lifetime of shed cells in the small intestine may be due to the high abundance of pancreatic enzymes that degrade biological material in this segment, or potentially due to the differential consistency of the luminal content between the small and large intestine.

Our study highlights an additional stage in the lifetime of intestinal cells that occurs outside of the tissue context. It will be important to use single cell approaches to study similar processes in the human gut in health and in diverse intestinal pathologies that may modulate shed cell dynamics. These include IBD, Celiac and colorectal cancer. Our work demonstrates that shed cells in the gut mirror pathological processes, and therefore could be utilized to probe gastrointestinal biology. Our approaches for combined analysis of transcriptomes of tissue and shed cells can be applied to study turnover dynamics in other mammalian tissues^37^.

## Methods

### Mice and sample collection

Mouse experiments were approved by the Institutional Animal Care and Use Committee of the Weizmann Institute of Science and were conducted in agreement with the institute guidelines. Male (C57BL/6JOlaHsd) mice aged 8-12 weeks, housed under regular 12h light-dark cycle, were used in our experiments, all mice were sacrificed by cervical dislocation. For control bulk RNA experiments, the whole intestine was taken out and divided into 5 parts; the small intestine from the end of the stomach to the cecum was divided to 3 equal parts of about 8 cm each. The large intestine from the end of the cecum to the rectum was divide to 2 equal length parts of about 2 cm each. For collection of fecal wash; each of the 5 segments was flushed with 700 ul of DPBS and was snap frozen in liquid nitrogen. After flushing, a small tissue fragment of 0.5 cm was taken from the distal part of each segment and snap frozen in liquid nitrogen. For the DSS colitis model, mice were treated with 2% (w/v) DSS (M.W. = 36,000–50,000 Da; MP Biomedicals) in their drinking water for 7 days followed by regular access to water for 3 days. Mice were monitored daily for weight loss (fig. 5a) and rectal bleeding signs. Colon fecal washes and tissue samples were taken at the 10th day of experiment. Since the colon was severely inflamed, it was taken as a whole and not split into two parts. For the antibiotics model, mice were given a combination of the following antibiotics for 4 weeks, vancomycin (1 g/l), ampicillin (1 g/l), kanamycin (1 g/l), and metronidazole (1 g/l) in their drinking water for 30 days^38–40^. All antibiotics were obtained from Sigma Aldrich. Colon fecal wash and tissue samples were taken at the 30th day of experiment.

### RNA isolation

For tissue samples – snap frozen tissues were thawed in 300 μl Tri-reagent and mechanically homogenized with bead beating, followed by a short centrifugation step to pull down beads and any tissue left-overs. For fecal washes – Tri-reagent was added to sample at a ratio of 3:1, samples were allowed to thaw on ice followed by thorough mixing. A first centrifugation step was used (1 minute, 18,000 rpm) to eliminate fecal solids. Following this, ethanol was added in a ratio of 1:1 to the supernatant from the previous step and continued according to the manufacturer instructions of Direct-zol mini (for tissues) and micro (for fecal wash) prep kit (ZYMO research, R2052).

### Bulk RNA sequencing of samples

RNA was processed by the mcSCRBseq protocol^18^ with minor modifications. For tissue RNA, RT reaction was applied on 20 ng of total RNA, for fecal wash RNA, RT reaction was started with one third of the total eluted volume with a final reaction volume of 20 μl (1× Maxima H Buffer, 1 mM dNTPs, 2 μM TSO* E5V6NEXT, 7.5% PEG8000, 20U Maxima H enzyme, 1 μl barcoded RT primer). Subsequent steps were applied as mentioned in the protocol. Library preparation was done using Nextera XT kit (Illumina) on 0.6 ng amplified cDNA. Library final concentration of 2nM was loaded on NextSeq 500/550 (Illumina) sequencing machine aiming at 20 M reads per sample with the following setting: Read1 – 16bp, Index1 – 8bp, Read2 – 66bp.

### Bioinformatics and computational analysis

Illumina output files were demultiplexed with bcl2fastq v.2.17. Resulting FASTQ files were analyzed using the zUMIs pipeline^41^. Reads were aligned with STAR (v.2.5.3a) to the GRCm38 genome (release 84; Ensembl) and exonic unique molecular identifier (UMI) counts were used for downstream analysis. Statistical analyses were performed with MATLAB R2021a. Mitochondrial genes and non-protein coding genes were removed from the analysis. Protein coding genes were extracted using the annotation in the Ensembl database (BioMart) for reference genome GRCm38 version 84, using the R package “biomaRt” (version 2.44.4). Gene expression for each sample was consequently normalized by the sum of the UMIs of the remaining genes that individually take up less than 5% of the sample sum. Samples with less than 100,000UMIs over the remaining genes were removed from the analyses. Clustering was performed with the MATLAB function clustergram over the Zscore-transformed expression matrix, using Spearman distances and included genes with maximal expression above 10^−4^ of summed UMIs. Differential gene expression was performed using Wilcoxon ranksum tests and Benjamini-Hochberg FDR corrections. Computational deconvolution was performed using cellanneal using signature tables obtained from a single cell atlas of small and large mouse intestines as follows (Supplementary Table 3). Villus bottom and top enterocytes were defined as the averages of the lower or upper 3 villus zones in Moor and colleagues^24^, respectively. Goblet cells and enteroendocrine cells signatures were extracted based on Haber and colleagues^19^. Immune cell signatures were extracted based on Biton and colleagues^42^. All dendritic cell subtypes were averaged and coarse-grained into one group, as were monocytes and macrophages. Large intestine signature file was extracted based on Širvinskas and colleagues aged and young colon single cell atlas^43^. Gene Set Enrichment Analysis (GSEA)^23^ was performed over the Hallmark gene sets.

### Mathematical model for cell shedding in the intestine

To estimate the contribution of each tissue segment to the fecal wash transcriptome (Fig. 1d) we used a multi-linear regression model:

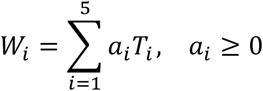

Where *W*_*i*_ is the i’th fecal wash segment, *T*_*i*_ is the i’th tissue segment, and *a*_*i*_ is a non-negative weight accounting for the contribution of the i’th tissue segment.

To estimate shed cell life-times, a simple model was formulated, in which the wash mRNA in each segment *W*_*i*_ is shed from the corresponding tissue segment *T*_*i*_ at rate *α*_*i*_, and degraded at rate *δ*_*i*_ :

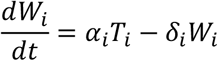

Where i=1…5 stands for duodenum, jejunum, ileum, proximal colon and distal colon respectively.

The equation was discretized, using the fact that the time axis in this problem corresponds to the spatial 1D axis along the intestinal tract:

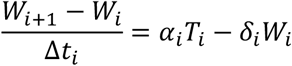

Where Δ*t*_*i*_ is the time it takes for the wash to move from the *i*’th segment to the (*i* + 1)’th segment. We took *t*_*i*_ to be *t*_1_, *t*_2_ *=* 1*h*, representing the transitions from duodenum to jejunum and from jejunum to ileum. *t*_3_, *t*_4_ *=* 3*h*, representing transitions from ileum to proximal colon and from proximal colon to distal colon following ^20–22^.

This leads to the final equation:

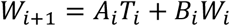

With *A*_*i*_ *= α*_*i*_ Δ*t*_*i*_, *B*_*i*_ *=* 1 *− δ*_*i*_ Δ*t*_*i*_.

The equation was fit for *i =* 1 … 4 by multi-linear regression with non-negative weights. The half-life time of the mRNA in each segment *τ* _*i*_ was computed from *B*_*i*_ by 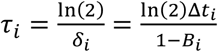. We excluded from the analysis genes that had expression level lower than 10^−4^ in all the tissue and wash samples. In Fig. 1f, the contribution of the current segment tissue and previous segment fecal wash were calculated as 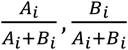 respectively. Violin plots in Fig. 1g and error bars in Fig. 1f were generated by bootstrapping the genes a hundred times (error bars represent standard errors of the mean).

### Single-cell isolation and sorting

Single cells were obtained from fecal washes of the entire mouse small intestine from four mice in total. Collected cells were filtered through 70μm filter and washed once with FACS buffer. Cells were stained with Hoechst (15 μg/ml) in the presence of reserpine (5 μM) for elimination of bacterial populations. Cells were sorted with SORP-FACSAriaII machine (BD Biosciences) using a 100 μm nozzle. Dead cells were excluded on the basis of propidium iodide (PI) incorporation. Sorted cells were high on hoechst and low on PI (Extended Data Fig. 2). Cells were sorted into 384-well cell capture plates containing 2 μl of lysis solution and barcoded poly(T) reverse-transcription (RT) primers for single-cell RNA-seq^26^. Barcoded single cell capture plates were prepared with a mantis automated liquid handling platform. Four empty wells were kept in each 384-well plate as a no-cell control during data analysis. Immediately after sorting, each plate was spun down to ensure cell immersion into the lysis solution, snap frozen on dry ice and stored at - 80°C until processed.

### MARS-Seq library preparation

Single cell libraries were prepared, as described in^26^. Briefly, mRNA from cells sorted into MARS-Seq capture plates were barcoded and converted into cDNA and pooled using an automated pipeline. The pooled sample was then linearly amplified by T7 in vitro transcription and the resulting RNA was fragmented and converted into sequencing ready library by tagging the samples with pool barcodes and Illumina i7 barcode sequences during ligation, reverse transcription and PCR. Each pool of cells was tested for library quality and concentration was assessed as described in^26^. Machine raw files were converted to fastq files using bcl2fastq package, to obtain the UMI counts, reads were aligned to the mouse reference genome (GRCm38.84) using zUMI packge ^41^ with the following flags that fit the barcode length and the library strandedness: -c 1-7, -m 8-15, -l 66, -B 1, -s 1, -p 16.

### scRNAseq data processing

For each single cell and for each gene we first subtracted the estimated background expression. Background was calculated for each 384-well plate separately, as the mean gene expression in the four empty wells. After subtraction, negative values were set to zero. Next, cells with total UMI counts lower than 300 or higher than 10,000 and total gene counts lower than 200 were removed. We used Seurat v4.1.0 package in R^44^ v4.1.2 to visualize and cluster the single cell RNAseq data (Fig. 3a, Fig. 4a). Gene expression measurements (UMIs per gene) were normalized for each cell by the summed UMI, multiplied by a scale factor 10,000, and then log-transformed. To avoid undesired sources of variation in gene expression, we used Seurat to regress out cell-cell variation driven by total number of UMIs. Cell clustering was based on PCA dimensionality reduction using the first 15 PCs, and a resolution value of 0.8. Cell type-specific markers were used to interpret the clusters cell types. The enterocytes cell cluster was determined based on average expression of Epcam higher than 10^−4^ and of and Muc2 lower than 10^−6^ fraction of UMI counts per cell. This cluster included 424 cells. This set of shed enterocytes was pooled with Epcam positive enterocytes isolated from intestinal tissue^28^ and re-ran Seurat. For each cell in the new UMAP (Fig. 4a) zonation score was calculated along the crypt villus axis, as the fractional sum of villus tip landmark genes as in^24^. To compare gene expression changes in fecal washes compared to tissue cells we sampled the tissue cells to match the zonation score distribution of the fecal wash enterocytes using the inverse transform sampling method^45^ (Fig. 4a).

### Single molecule Fluorescence in-situ Hybridization (smFISH)

Fecal wash cells from the entire small intestine were centrifuged at 500g for 5 minutes, pellet of cells was fixed with pre-chilled 4% FA for 15 minutes in 4°C followed by 2hr incubation 70% EtOH. SmFISH experiments were performed on suspended cells. Probe libraries were designed using the Stellaris FISH Probe Designer Software (Biosearch Technologies, Inc., Petaluma, CA). Hybridization was performed according to published protocol^46^ with modifications for cell suspension. Briefly, cells were washed with 2× SSC (Ambion AM9765) for 5min. Cells were incubated in hybridization buffer (10% Dextran sulfate Sigma D8906, 20% Formamide, 1mg/ml E.coli tRNA Sigma R1753, 2× SSC, 0.02% BSA Ambion AM2616, 2 mM Vanadylribonucleoside complex NEB S1402S) mixed with probes overnight in a 30°C incubator. smFISH probe libraries were coupled to Cy5, TMR or Alexa594. Supplementary Table 5 includes a list of all the probe sequences used in this study. After the hybridization, cells were washed with wash buffer containing 50ng/ml DAPI for 30 min at 30°C. DAPI (Sigma-Aldrich, D9542) was used for nuclear staining. All images were performed on a Nikon-Ti-E inverted fluorescence microscope using the NIS element software AR 5.11.01.

### Intravital imaging by two-photon laser scanning microscopy (TPLSM)

Mice were anesthetized with a mixture of 50 mg ketamine, 15 mg xylazine and 2.5 mg acepromazine per kg of body weight. Mice were mounted in a custom-designed heated stage, kept at 37°C and supplemented with half dose of the anesthesia above every hour. A 1 cm incision was made in the abdominal skin and the midline and proximal intestinal tissue was externalized and a 15mm incision using a cautery was done to expose the luminal surface. Tissue was immobilized to a custom-made heated chamber using tissue adhesive (Vetbond, M3). Imaging area was hydrated with PBS using silica gel and a glass cover slip was placed at the region of interest. Intravital imaging was performed using a water-dipping objective (Zeiss 20X 1.05 NA plan objective). Zeiss LSM 880 upright microscope fitted with Coherent Chameleon Vision laser was used. Images were acquired with a femtosecond-pulsed two-photon laser tuned to 900 nm. The microscope was fitted with a filter cube containing 565 LPXR to split the emission to a PMT detector (with a 579-631 nm filter) and to an additional 505 LPXR mirror to further split the emission to 2 GaAsp detectors (with a 500-550nm filter for GFP fluorescence). Images were acquired as 33μm z-stacks with 3 μm steps between each z-plane (12 frames) at minimal interval for 30-60 min. The zoom was set to 1.5, and images were acquired at 512×512 x-y resolution. CX3CR1^GFP^ mice were intravenously injected with 4ul Hoechst 33342 (Thermo Fisher Scientific) in 100μl PBS 10 min prior imaging. For TNF-α treated mice, mice were injected with 10μg TNF-α intraperitoneal in PBS 20 min prior to the surgery.

## Acknowledgements

We thank Hagit Shapiro and Eran Elinav for help with experimental procedures of mice perturbations. S.I. is supported by the Wolfson Family Charitable Trust, the Edmond de Rothschild Foundations, the Fannie Sherr Fund, the Dr. Beth Rom-Rymer Stem Cell Research Fund<u>, the Helen and Martin Kimmel Institute for Stem Cell Research</u>, the Minerva Stiftung grant, the Israel Science Foundation grants No. 1486/16 and No. 3663/21, the European Research Council (ERC) under the European Union’s Horizon 2020 research and innovation program grant No. 768956, the Chan Zuckerberg Initiative grant No. CZF2019-002434, the Bert L. and N. Kuggie Vallee Foundation and the Howard Hughes Medical Institute (HHMI) international research scholar award.

## Data availability

All data generated in this study are available at the Zenodo repository: https://doi.org/10.5281/zenodo.7137319

## Extended Data

**Extended Data Fig. 1.**
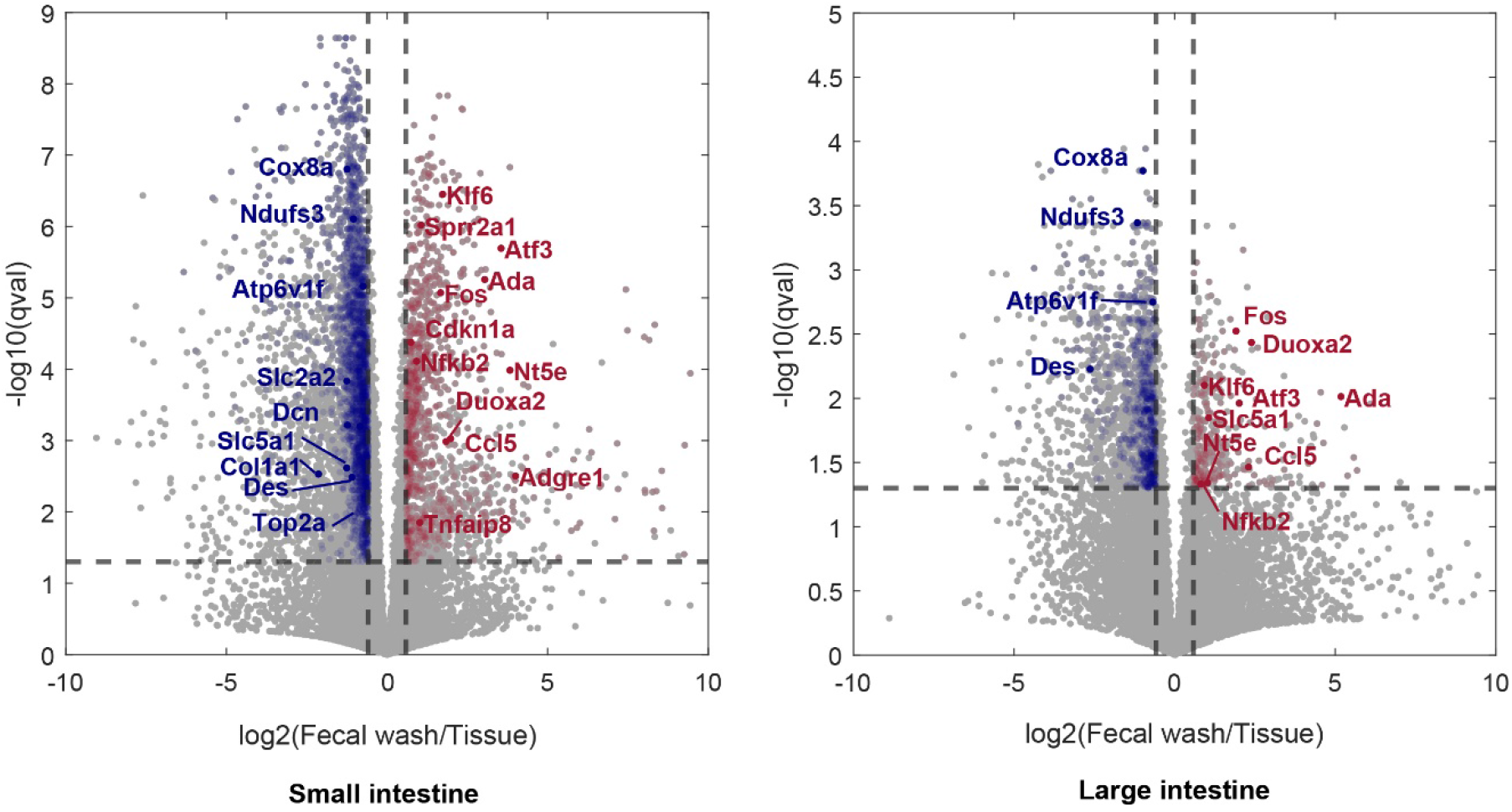
Volcano plot showing differentially expressed genes between fecal washes and tissues in small (left) or large (right) intestine, each dot is a gene, x-axis is the log2-ratio of expression between fecal wash and tissue, y axis is –log10 (q value), where p value is computed using Wilcoxon rank-sum tests and q-values computed using Benjamini-Hochberg FDR correction. Genes with corresponding q-values below 0.05 and fold change greater than 1.5 are marked in red or blue for fecal wash or tissue elevated genes respectively. Names of representative up-regulated genes are shown in red or blue for fecal wash or tissue respectively.

**Extended Data Fig. 2.**
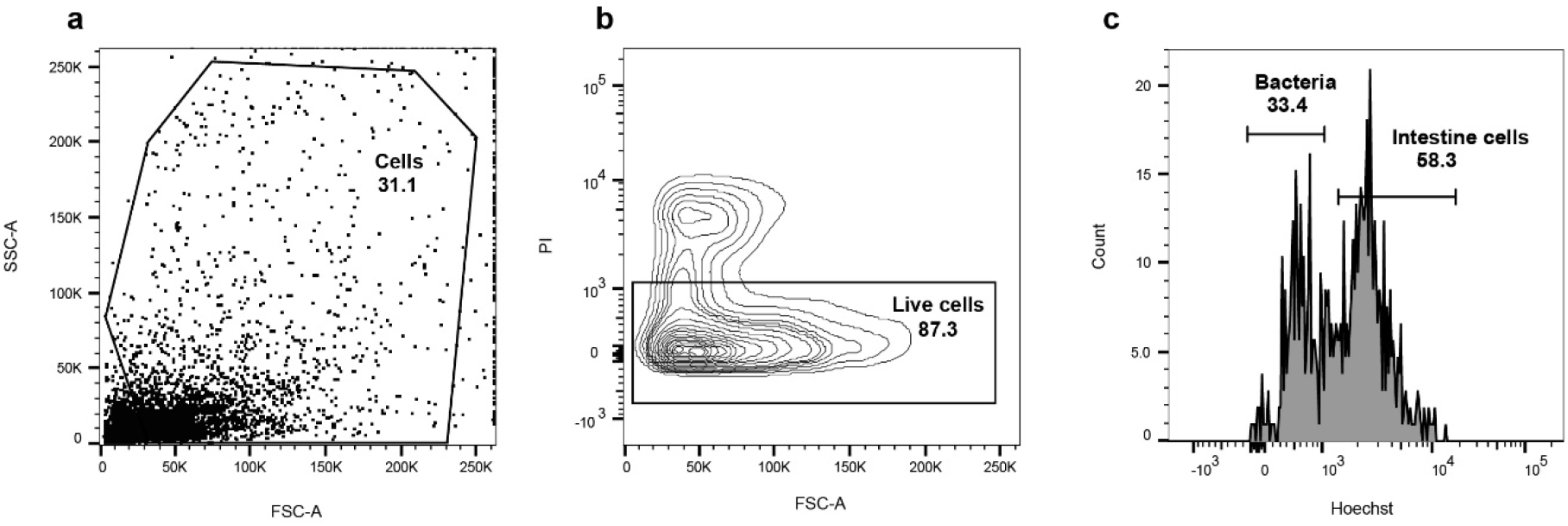
Sorting strategy for isolation of mouse intestinal cells from fecal washes. **a**. FSC-A and SSC-A are used to select cells based on size. **b**. PI staining is used to gate out dead cells. C. Hoechst fluorescense is used to select eukaryotic cells while avoiding low DNA amount of bacteria cells. Numbers in A-C represent the percent of gated cells.

**Extended Data Fig. 3.**
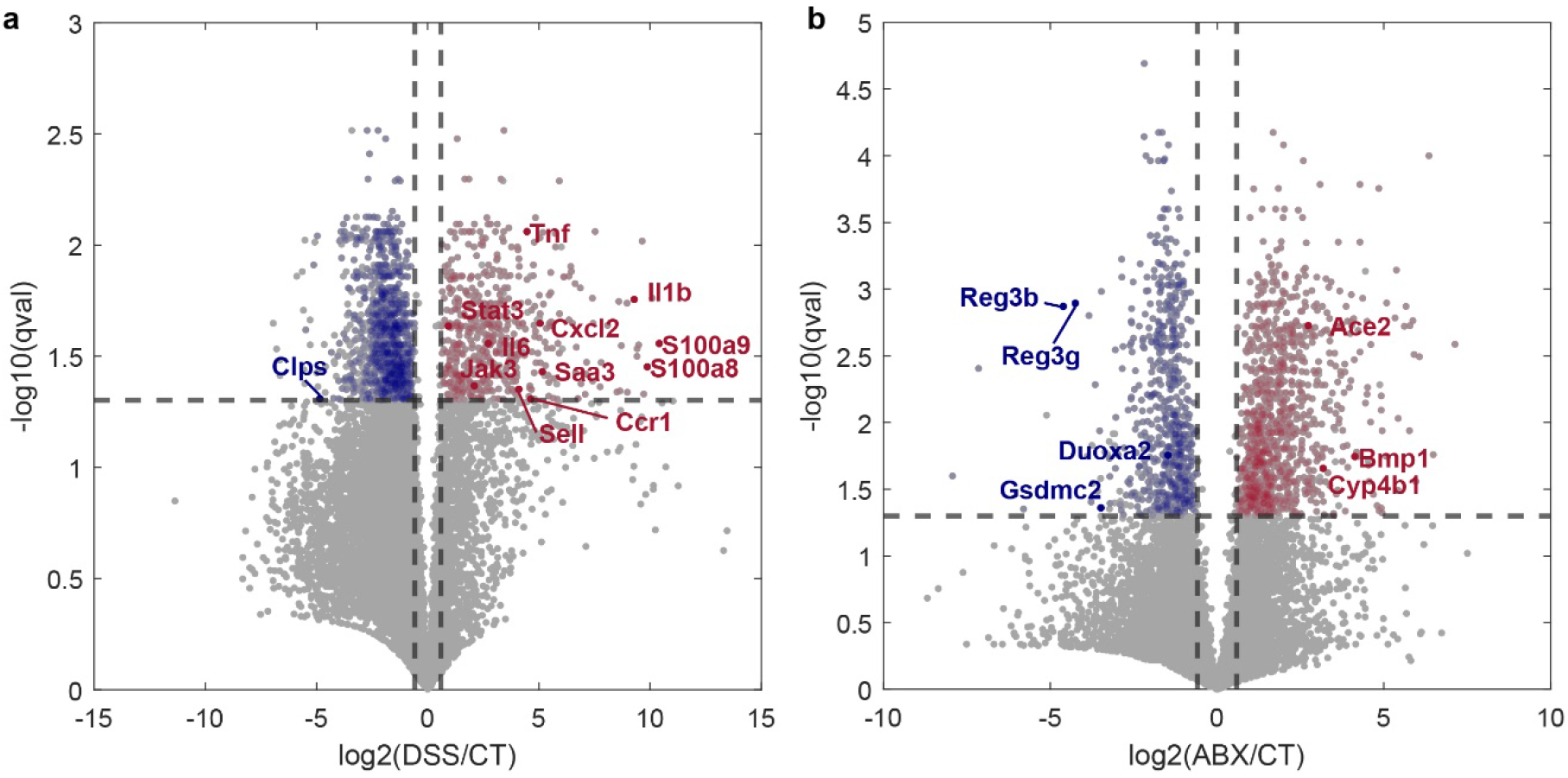
Volcano plot showing differentially expressed genes between large intestine fecal wash in control (CT) and DSS (a) or ABX and CT (b), each dot is a gene, x-axis is the log2-ratio of expression between fecal wash and tissue, y axis is –log10 (q value), where p value is computed using Wilcoxon rank-sum tests and q-values computed using Benjamini-Hochberg FDR correction. Genes with corresponding q-values below 0.05 and fold change garter than 1.5 are marked in red or blue for treatment or control elevated genes respectively. Names of representative up-regulated genes are shown in red or blue for treatment or control respectively.

